# Metaplastic priming enables non-ionotropic NMDA receptor-mediated synaptic depotentiation in the hippocampus

**DOI:** 10.1101/2025.02.28.640846

**Authors:** Quinn Pauli, Juliet Arsenault, Samuel W. Fung, Amy J. Ramsey, Robert P. Bonin

## Abstract

The reversal of learning-induced synaptic potentiation through depotentiation is thought to underlie forgetting and can be influenced by prior synaptic activity. Here, we evaluated how such metaplastic alterations manifest at the synaptic level. In hippocampal slices obtained from male and female mice, we artificially induced long-term potentiation (LTP) using either a temporally spaced or compressed stimulation pattern. Using a combination of electrophysiology and protein quantification approaches, we found divergent molecular pathways recruited during depotentiation of spaced and compressed LTP. Depotentiation of both forms of LTP required glutamatergic activation of the NMDA receptor (NMDAR). However, only depotentiation of spaced LTP required ionotropic NMDAR signaling, while ion flux-independent, or non-ionotropic, NMDAR signaling was necessary and sufficient for depotentiation of compressed LTP. Downstream of NMDAR signaling, AMPA receptor phosphorylation was also differentially modified during depotentiation of spaced and compressed LTP. Finally, we found that spaced but not compressed depotentiation required synaptic Arc. Together, our results reveal that the temporal pattern of prior LTP induction exerts a metaplastic influence on the molecular pathways recruited during the induction and expression of depotentiation. Our findings have important implications for the regulation of both physiological and pathological forgetting.

**Significance statement:** Synaptic depotentiation, the reversal of learning-associated synaptic potentiation, is an important mechanism of forgetting. This study uncovers how prior synaptic activity modifies the molecular mechanisms underlying depotentiation in the hippocampus in a metaplastic manner. We reveal that the mechanisms of NMDA-receptor depotentiation depend on the temporal spacing of long-term potentiation (LTP) induction. Specifically, we show that non-ionotropic NMDA receptor signaling is necessary and sufficient for the depotentiation of LTP induced using temporally compressed but not spaced patterned activity. Further, depotentiation of spaced and compressed LTP are associated with distinct downstream signaling pathways and AMPA receptor phosphorylation states during depotentiation. Our results illuminate fundamental mechanisms that govern plasticity associated with forgetting, with implications for memory preservation in disease states.

## Introduction

Plasticity at the network, synaptic and molecular levels of the brain give rise to adaptive learning and forgetting processes. Activity-dependent tuning of synaptic transmission among cells that comprise a memory (engram cells) is thought to modulate memory accessibility and longevity (Han et al., 2021). Correspondingly, the history of activity at a synapse can affect subsequent plasticity, referred to as metaplasticity (Abraham and Bear, 1996). For example, although they can be induced using similar stimuli, long-term depression (LTD) at naïve synapses and depotentiation (DEP) of synapses that have undergone long-term potentiation (LTP) are distinct processes (Huang and Hsu, 2001). LTD and DEP may also serve different physiological functions, where DEP of synapses that have undergone learning-induced potentiation may be particularly relevant in forgetting (Ge et al., 2019; Moreno, 2021). Besides the presence of LTP, the specific type of LTP induced can also regulate DEP. LTP induced with repeated theta burst stimulations (TBS) with increasing temporal spacing correlates with more stable LTP (Buschler et al., 2012) and progressively lower DEP (Woo and Nguyen, 2003; Park et al., 2019). Different stabilizing mechanisms are set during LTP induced with a temporally spaced or compressed TBS pattern (Park et al., 2016; Park et al., 2021). However, a major unknown is how the metaplastic effects of LTP induction manifest during DEP.

Synaptic plasticity in the hippocampus is expressed primarily through postsynaptic alterations in α-amino-3-hydroxy-5-methyl-4-isoxazolepropionic acid receptor (AMPAR)-mediated signaling. AMPARs have also been implicated in metaplasticity, where distinct sites on the GluA1 subunit are specifically dephosphorylated during LTD and DEP (Lee et al., 2000). Different AMPAR subunits may also prevent or promote forgetting. For instance, synaptic GluA2 is elevated in the hippocampus following strong, but not weak, inhibitory avoidance training associated with long-lasting memory (Dong et al., 2015). Changes in AMPAR function and expression are initiated by signaling from synaptic N-methyl-D-aspartate receptors (NMDARs).

NMDARs mediate memory processes as well as bidirectional forms of hippocampal synaptic plasticity including DEP (Huang et al., 2001; Luscher and Malenka, 2012). Classically, NMDAR-mediated Ca^2+^ influx initiated by concurrent glutamate and co-agonist (glycine/D-serine) binding and postsynaptic depolarization has been viewed as a major plasticity trigger. A growing body of evidence has demonstrated that glutamate receptors, including the NMDAR, can also signal in an ion flux-independent manner. Glutamate binding alone can cause conformational changes in the C tail of the GluN1 receptor subunit, triggering signaling cascades that cause synaptic weakening (Aow et al., 2015; Dore et al., 2015). Non-ionotropic NMDAR (NI-NMDAR) signaling has specifically been implicated in LTD, spine shrinkage and certain disease states (Kessels et al., 2013; Nabavi et al., 2013; Stein et al., 2015). It is important to note that in contrast, others have shown a critical role for ionotropic NMDAR signaling during LTD (Babiec et al., 2014). Additionally, the type of NMDAR signaling involved in DEP remains unclear (Latif-Hernandez et al., 2016; Ge et al., 2019). Recently, our lab has shown that reactivation-dependent weakening of potentiated, but not naïve, spinal pathways requires NI-NMDAR signaling (Zhang et al., 2023), suggesting that prior synaptic activity may determine the effects of NI-NMDAR signaling.

Given the evidence that prior synaptic activity can regulate NI-NMDAR signaling and AMPAR expression, we evaluated the influence of LTP induction on the mechanisms recruited during DEP in the hippocampus.

## Materials and Methods

### Animals

All animal care and experimental procedures were reviewed and approved by the Animal Ethics and Compliance Program at the University of Toronto and conducted in accordance with the Canadian Council on Animal Care (CACC) guidelines. Mice were kept on a 14/10h light-dark cycle in groups of one to four mice per cage with food and water provided *ad libitum*. The ARC^tm1.1(cre/ERT2)Luo^ (*Arc^CreER^*) knock-in mouse line (JAX:021881, The Jackson Laboratory, Bar Harbor, Maine, United States) contains a Cre recombinase insertion (CreER^T2^) and replaces the endogenous 3’ untranslated region (3’ UTR) of the *Arc* loci. This mutation specifically disrupts dendritic Arc mRNA trafficking (Kobayashi et al., 2005). The *Arc^CreER^* mice were bred on a C57Bl/6J background and were homozygous for this mutation.

### Preparation of hippocampal slices

Field excitatory postsynaptic potential (fEPSP) recordings were performed at Schaffer collateral-CA1 synapses in acute parasagittal hippocampal slices. 6-12-week-old male and female wildtype (WT) C57BL/6J (The Jackson Laboratory) or C57BL/6NCrl (Charles River Laboratories, Wilmington, Massachusetts, United States) and *Arc^CreER^* mice were first anesthetized with 2.5% Avertin (Tribromoethanol) and euthanized by trans-cardiac perfusion with an ice cold (2-4^0^C) sucrose-based solution containing (in mM): 50 Sucrose, 92 NaCl, 15 Glucose, 5 KCl, 1.4 NaH_2_PO_4_, 26 NaHCO_3_, 0.5 CaCl_2_, 7 MgCl_2_6H_2_O and 1 Kynurenic acid (osmolality 322-330 mOsm, bubbled with 95% O_2_ and 5% CO_2_). The brain was then removed and allowed to rest for 2 minutes in ice cold artificial cerebrospinal fluid (ACSF) containing (in mM): 124 NaCl, 10 glucose, 26 NaHCO_3_, 3 KCl, 1.4 NaH_2_PO4, 1 MgSO_4_ and 2 CaCl_2_ (osmolality 300-310 mOsm, bubbled with 95% O_2_ and 5% CO_2_). The cerebellum and olfactory bulb were removed, and the brain was hemisected for slicing. Parasagittal dorsal hippocampal slices (400 µm) were cut with a Leica VT1200 S vibratome (Leica Biosystems, Deer Park, Illinois, United States) in ice cold ACSF, and the CA3 region was carefully removed from each slice. Slices were transferred to a recovery chamber and maintained in 28^0^C ACSF bubbled with 95% O_2_ and 5% CO_2_ before use, for a minimum of 1.5 hours before recordings were made. Slices were prepared within 5 hours of the start of animals’ daylight cycle. All chemicals for slice preparation and electrophysiology recordings were purchased from Sigma Aldrich Canada (Oakville, Ontario, Canada). Hippocampal slice preparation protocol also available on protocols.io (dx.doi.org/10.17504/protocols.io.eq2ly6j9egx9/v1).

### Slice electrophysiology

Extracellular electrophysiology was performed on hippocampal slices transferred to a submerged-type recording chamber and continuously perfused at a rate of 1.5 mL/min with oxygenated ACSF maintained at 28.5^0^C (± 0.5^0^C) with an in-line solution heater and heated platform controlled by a dual channel temperature controller (Warner Instruments, Hamden, Connecticut, United States). ACSF was also pre-heated with a 4 channel BubbleStop (AutoMate Scientific, Berkeley, California, United States). An ACSF-filled borosilicate glass pipette (1-3 MΩ) was placed on the surface of the stratum radiatum in CA1. A bipolar platinum-iridium stimulating electrode (cat nr. #30250, FHC, Bowdoin, Maine, United States) controlled by a constant current stimulus isolator (A365, World Precision Instruments, Sarasota, Florida, United States) (0.1 ms pulse width) was used to evoke field excitatory postsynaptic potentials (fEPSPs) at the Schaffer collateral-commissural pathway. Test pulses were evoked every 30s (0.033 Hz). Stimulus input/output curves for fEPSPs were generated using 10-28 µA pulses in increments of 2 µA while monitoring evoked responses. To induce LTP, the stimulus strength was adjusted to maintain the fEPSP peak amplitude at half-maximal responses according to the input/output curve. A stable baseline lasting at least 30 minutes was recorded before LTP induction. LTP was induced at the same basal stimulus intensity using either a spaced or compressed theta burst pattern (sTBS, cTBS), consisting of five bursts at 5Hz, with each burst composed of five pulses at 100Hz and each train repeated 3 times spaced apart by either 10 minutes (sTBS) or 10 seconds (cTBS) (as in (Park et al., 2016)). 30 minutes after the last TBS train, DEP was induced using a low frequency stimulation (2Hz, 10 minutes LFS) as in (Park et al., 2019). In experiments with drug application, drugs were applied beginning 10-15 minutes after the last train of TBS and washed out 5-15 minutes after the LFS as indicated in each figure. fEPSPs were monitored for at least 35 minutes after the induction of DEP. Paired-pulse ratio (PPR) responses to two impulses delivered at an inter-stimulus interval (ISI) of 50, 100, 150, 200, 250 or 500 ms were recorded. For experiments where PPR responses were collected at baseline, LTP and post-LFS to monitor presynaptic changes, an ISI of 50 ms only was used. Signals were amplified using a Multiclamp 700A amplifier (Axon Instruments, Scottsdale, Arizona, United States) and were low pass filtered at 2.4 kHz and digitized at a sampling rate of 20 kHz with an Axon Digidata 1550B (Axon Instruments). Recordings were monitored and analyzed using pClamp v10.7 (Axon Instruments).

### Electrophysiology analysis

The evoked fEPSP descending slope (mV/ms) was measured from the fiber volley to the peak (10%-90%) and expressed as a percentage of the mean baseline response. The percent DEP was calculated by subtracting a five-minute fEPSP average 20-25 minutes (sDEP), or 25-30 (cDEP) minutes after administration of the LFS (DEP) from a five-minute fEPSP average 10-15 minutes after the final TBS administration (LTP). The timepoints were chosen to ensure that responses had stabilized after plasticity induction and drug washout. To probe for presynaptic changes that occur specifically during DEP, the percent change in PPR (50 ms ISI) was calculated between LTP and DEP. Similar to percent DEP, the PPR percent change was calculated by subtracting the PPR obtained after LFS administration (DEP) from the PPR obtained after LTP induction, normalized to the baseline PPR. fEPSP peak amplitude (mV) was used to obtain input/output curves and PPRs. Representative traces were smoothed with consistent parameters, and stimulus artifacts were excluded for clarity.

### Drugs

Drugs were prepared as stock solutions (50-100 mM) in distilled water or dimethyl sulfoxide (DMSO) (L-689,560 only) and then diluted in ACSF. The concentration of DMSO was 0.01% in the final solution. The glutamate-site specific NMDAR antagonist D-(-)2-Amino-5-phosphonopentanoic acid (APV) was obtained from HelloBio (Bristol, United Kingdom), and the glycine-site specific NMDAR antagonist L-689,560 (L689) was obtained from Tocris Bioscience (Biotechne, Minneapolis, Minnesota, United States). 7-Chlorokynurenic acid (7-CK), a different glycine site antagonist was obtained from Abcam (Cambridge, United Kingdom).

### NMDAR mimetic Tat peptides

As previously described (Zhang et al., 2023), the cell permeable NMDAR mimetic peptide C1.1. corresponds to amino acids 864-877 of GluN1 (DRKSGRAEPDPKKK). C1.1 and its associated scrambled control, C1.1Scr (EPAKDDGRRKPSKK), were synthesized with a Tat sequence (YGRKKRRQRRR) at the C-terminal to facilitate cell membrane internalization. Both peptides were synthesized by GenScript (Piscataway, New Jersey, United States).

### Immunofluorescence

GluA1 and GluA2 subunit expression was quantified in hippocampal slices after sLTP, cLTP, sDEP or cDEP induction using immunofluorescence. After electrophysiological recordings were terminated (30-40 minutes after plasticity induction), slices were drop fixed in 4% paraformaldehyde in phosphate-buffered saline (PBS) for 2 hours (room temperature). Control naïve slices were incubated in 28^0^C ACSF for a minimum of 2 hours prior to fixation. Slices were then washed three times with PBS and transferred to 30% sucrose in PBS (w/v). After cryopreservation for 1 week in 30% sucrose solution, slices were stored at −80^0^C in cryoprotectant. Next, slices were cryosectioned at 20 µm and mounted onto microscope slides. Sections were washed 5×5 minutes in 0.1% Triton-X in PBS (0.1% PBS-T). Non-specific binding was blocked using 10% horse serum in 0.1% PBS-T for 1 hour. Samples were then incubated with primary antibodies in blocking buffer overnight at 4^0^C and then washed 5×5 minutes in PBS-T. Next, they were incubated with secondary antibodies for 2 hours at room temperature in blocking buffer and then washed 5×5 minutes in PBS-T. Lastly, they were incubated with DAPI (1:500, Abcam, cat nr # ab228549) for 10 minutes in PBS at room temperature and washed 2×5 minutes in PBS. Primary antibodies used include anti-GluA1 (1:400, guinea pig host, Alomone labs, Jerusalem, Israel, cat nr #AGC-004-GP, RRID: AB_2340961), anti-GluA2 (1:1000, rabbit host, Abcam, cat nr #ab206293, RRID: AB_2800401), and NeuN (1:1000, mouse host, Millipore-Sigma, Burlington, MA, United States, cat nr #MAB377, RRID: AB_2298772). Secondary antibodies used include anti-mouse Alexa 647 (1:500, Jackson ImmunoResearch Labs, West Grove, Pennsylvania, United States, cat nr #115-606-003, RRID: AB_2338921), anti-guinea pig Alexa-488 (1:500, Jackson ImmunoResearch Labs, cat nr #706-545-148, RRID: AB_2340472), and anti-rabbit Alexa-594 (1:500, Thermo Fisher Scientific, Waltham, Massachusetts, United States, cat nr #A-11037, RRID: AB_2534095). Slides for imaging were mounted with Prolong Glass Antifade (Thermo Fisher, P36984) and sealed with clear nail polish to prevent tissue from drying out. Protocol also available on protocols.io (dx.doi.org/10.17504/protocols.io.4r3l2923qv1y/v1).

### Confocal microscopy and image quantification

Images were acquired using a Leica SP8 confocal microscope (frame sequential scan), using a 20x objective to obtain z stack images (6 µm, maximum intensity projection) of the CA1 region. Identical gain and laser power were used to capture all images. Image contrast adjustments were linear and uniform across all representative slices, for display purposes only. For slices collected following electrophysiology recordings, sections with visible electrode marks to identify the recording plane were chosen for imaging whenever possible. Average fluorescence intensity (mean grey value) and the intensity profile in the GluA1 and GluA2 channels for quantification was calculated using ImageJ software in the stratum radiatum (SR), with a region of interest defined for each slice to account for slight variations in morphology. Contour lines were drawn around the SR based on differences in GluA1 and GluA2 expression demarcating the SR and the stratum lacunosum-moleculare (SLM), and below the pyramidal cell bodies marked using NeuN. Mean fluorescence intensity comparisons were made to quantify relative amounts of AMPAR subunit expression across LTP and DEP groups (GluA1/GluA2) in the SR. To account for differences in fluorescence that may arise due to experimental variation, the values for each experimental slice (sLTP, cLTP, sDEP, cDEP) were normalized to a naïve slice stained and imaged within the same batch of experimental slices. A maximum of 4 experimental slices were normalized to a single naïve slice. To quantify the spatial distribution of GluA1 and GluA2, average intensity values as a function of distance along the SR were obtained using the plot profile function along a rectangular region enclosing the isolated SR. The distance along the SR was normalized to a 0-1 scale to account for morphological variation among slices. Intensity values were normalized to the maximum intensity value. Background subtraction was performed using the mean grey value of a region outside of each slice prior to all quantitative comparisons.

### Western blotting and quantification

Hippocampal slices were collected after electrophysiology recordings and flash frozen at −80^0^C. Naïve slices were collected and flash frozen after resting in 28^0^C ACSF for at least 2 hours. The hippocampus (including the dentate gyrus) was microdissected on dry ice, homogenized in 100µL ice cold lysis buffer containing 50 mM Tris-HCl (pH 7.5), 150 mM NaCl, 5 mM EDTA, 5 mM EGTA (pH 8.0), 20 mM NaF, 1% Triton X-100, phosphatase inhibitor cocktail II and III (Millipore-Sigma) and EDTA-free protease inhibitor cocktail table (Roche Diagnostics, Indianapolis, Indiana, United States) using a Dounce homogenizer. Debris was removed by centrifugation at 12,000 g for 15 minutes at 4^0^C. The supernatant was transferred to a new tube and kept on ice for further processing. The amount of total protein was assayed for each sample (Pierce bicinchoninic acid). Whole lysate samples were mixed in 2 x Laemmli buffer (65.8 mM Tris-HCl, pH 6.8, 2.1% sodium dodecyl sulphate (SDS), 26.3% (w/v) glycerol, 0.01% bromophenol blue and 5% β-mercaptoethanol; Bio-Rad Laboratories, Hercules, California) and boiled at 95^0^C for 5 minutes to prepare for SDS-PAGE gel electrophoresis. 5 µg samples were loaded into 10% polyacrylamide gels, and the separated proteins were transferred to nitrocellulose membranes at 200mA overnight (Mini-PROTEAN Tetra Cell, Mini Trans-Blot Module, and PowerPac Basic Power Supply, BioRad Laboratories). Membranes were briefly stained with 0.1% Ponceau and then incubated in blocking buffer (5% milk in Tris-buffered saline with Tween20 (TBS-T; 20 mM Tris, 150 mM NaCl, 0.1% Tween20)) for 1 hour at room temperature. The blotted proteins were then probed with moloclonal anti-GluA1 (1:1000, Cell Signaling Technology (CST), Danvers, Massachusetts, United States, cat nr #13185, RRID: AB_2732897), anti-GluA2 (1:1000, CST, cat nr #13607, RRID: AB_2650557), anit-GluA1-Ser831 (1:1000, CST, 75574, RRID: AB_2799873), anti-GluA1-Ser845 (1:1000, CST, 8084, RRID: AB_10860773) or anti-GAPDH (1:12000, CST, cat nr #2118, RRID: AB_561053) overnight at 4^0^C in 5% bovine serum albumin (BSA) in TBS-T. Following primary antibody incubation, membranes were washed in 1 X TBS-T (3 x 10 minutes). The blots were then probed with HRP-conjugated secondary antibodies at 1:5000 (goat antirabbit, CST, cat nr #7074, RRID: AB_2099233) for 2 hours at room temperature in 5% BSA. The proteins were visualized by incubating membranes in ECL developing solutions (Clarity Western ECL Substrate, cat nr. #1705061, BioRad) mixed in equal volumes for 1 minute followed by imaging with a CCD camera-based imager (ChemiDoc Imaging System, BioRad). Exposure times were adjusted for each antibody and blot to ensure bands were not overexposed and were quantifiable. Linear contrast adjustments to images were kept consistent within an image to enhance band visibility, without distorting bands. Band intensity was quantified using integrated density of a region of interest of consistent area for each blot, with a consistent background region selected within each lane. The background was subtracted for each band before normalization, using an area within the same lane above or below the band. Within each blot, GluA1, GluA2 and pGluA1 levels for each slice were first normalized to the corresponding amount of GAPDH (loading control) and then normalized to a naïve slice to account for experimental variation across replicates and to enable comparisons across blots. Phosphorylated GluA1 at S831 and S845 was then normalized to total GluA1. sDEP and cDEP conditions were run separately and compared to one another only after normalization. All proteins (GluA1, GluA2, pGluA1 S831 and pGluA1 S845) were probed with replicates from the same tissue sample. Quantification was performed by two experimenters and obtained values were averaged.

### Tissue Fractionation, RNA Isolation, and RT-qPCR

Our synaptosome preparation was adapted from previous publications to favour the isolation of the postsynaptic membrane (Hafner et al., 2019; Zhang et al., 2023). The hippocampus was removed and stored in −80°C until processing. A single hippocampus was homogenized in a microtube homogenizer (Bel-art) in 200 µl of ice-cold homogenizing buffer (0.32 M sucrose and 5 mM HEPES (pH 7.4)), supplemented with a Murine RNAse inhibitor (M0314S, New England Biolabs, Ipswich, Massachusetts, United States) at the manufacturer’s recommended concentration. An aliquot of the lysate was saved as ‘total RNA’. The remainder was centrifuged (1000g for 10 minutes at 4°C). The supernatant (S1) was further centrifuged (12,000g for 12 minutes at 4°C) to obtain the synaptosome-enriched pellet (P2) from the supernatant (S2). The P2 pellet was stored in TRI-REAGENT (TRI118, BioShop Canada, Burlington, Ontario, Canada) at −80°C until RNA isolation.

To isolate RNA from the cytoplasmic fraction, an RNA precipitating solution (0.15 M sodium acetate in 100% ethanol) was added to the supernatant (S2) and incubated in −20°C for 1 hour (Gagnon et al., 2014). The sample was vortexed for 30 seconds and centrifuged (18,000g for 15 minutes at 4°C). The supernatant was discarded, washed by vortexing briefly in ice-cold 70% ethanol, and centrifuged (18,000g for 5 minutes at 4°C). The supernatant was discarded and the resulting cytoplasmic-enriched pellet was partially dried. The cytoplasmic pellet was stored in TRI-REAGENT −80°C until RNA isolation. Total, cytoplasmic, and synaptosomal samples were processed for RNA using TRI-REAGENT according to the manufacturer’s guidelines. RNA was resuspended in TE Buffer (0.1 M Tris-Cl and 0.01 M EDTA, pH 8). RT-qPCR was performed using a one-step kit (E3005S, New England Biolabs) on total, cytoplasmic, and synaptosomal RNA. Fold change was calculated using the ΔΔCT method and normalized to wild-type samples. Primers obtained from Integrated DNA Technologies (Coralville, Iowa, United States) (Arc-F: TGTTGACCGAAGTGTCCAAG, Arc-R: AAGTTGTTCTCCAGCTTGCC, GAPDH-F: GGCAAATTCAACGGCACAGT, GAPDH-R: GGGTCTCGCTCCTGGAAGAT).

### Experimental Design and Statistical Analysis

All electrophysiology treatment groups were interleaved with control experiments, with slices from at least five different mice. A maximum of 2 slices from the same animal were used in a treatment condition, where slices from the same animal were most commonly used in two different conditions within the same dataset. Datapoints and n values for each figure represent individual slices. Immunoblotting and immunofluorescence experiments were performed with slices from at least three mice, with 1-2 technical replicates averaged together. In all figures, results are expressed as means ± standard error of the mean (SEM). Dotted lines in all representative traces indicate the average baseline response used for normalization of fEPSPs. All scalebars for representative traces are 5 ms x 0.2 mV. For electrophysiology experiments, the percent DEP was calculated by subtracting a five-minute fEPSP average 20-30 minutes after administration of the LFS (after drug washout) from a five-minute fEPSP average 10-15 minutes after the final TBS administration. Statistical analysis was performed using GraphPad Prism v10. Treatment groups were compared using a two-tailed paired or unpaired Student’s t test or a one- or two-way ordinary or repeated-measures ANOVA as indicated in figure captures. For repeated measures ANOVAs, sphericity was not assumed, and the method of Geisser and Greenhouse was used to correct for violations of the assumption. Adjustments were made for multiple comparisons using the Holm-Šídák correction. When values were missing due to unequal sample sizes in a repeated measures ANOVA, we analyzed the data instead by fitting a mixed model which in the absence of missing values, gives the same p values and multiple comparisons tests as repeated measures ANOVA. As such, in the presence of missing values we interpreted the results like a repeated measures ANOVA. For all experiments, statistical significance was set at p ≤ 0.05 (*p ≤ 0.05, **p ≤ 0.01, ***p ≤ 0.001), and reported p values were multiplicity adjusted where appropriate to account for multiple comparisons.

## Results

### NMDAR glutamate- and glycine-site antagonists differentially inhibit cDEP and sDEP

To begin to assess the possible metaplastic constraints on the mechanisms recruited during DEP, we pharmacologically isolated NI-NMDAR signaling in ex vivo dorsal hippocampal slices from 6-12-week-old C57Bl/6 mice. Field excitatory postsynaptic potentials (fEPSPs) recorded in the CA1 region were evoked by stimulating Schaffer collateral inputs every 30 seconds. After establishing a stable baseline at half-maximal responses, LTP was induced with 3 TBS repetitions spaced apart by either 10 minutes (sLTP) or 10 seconds (cLTP). DEP was triggered using a 2Hz LFS (1200 pulses) 30 minutes after administration of the final TBS. We observed that DEP of both cLTP (cDEP) and sLTP (sDEP) were readily induced in slices prepared from male mice (Fig. 1A). Next, to distinguish between a requirement for NMDAR activation and ionotropic receptor functioning, we bath-applied the glycine site-specific antagonist 7-Chlorokynurenic acid (7-CK) (100 µM) previously shown to block synaptic and extrasynaptic NMDAR currents, promoting NI-NMDAR signaling (Nabavi et al., 2013; Stein et al., 2015). Isolating NI-NMDAR signaling with 7-CK blocked sDEP on average (Fig. 1B, right), consistent with a requirement for ionotropic NMDAR signaling. In contrast, cDEP persisted in the presence of 7-CK (Fig. 1B, left), suggesting that NI-NMDAR signaling may be sufficient to induce this type of DEP. We additionally confirmed that both cDEP and sDEP required glutamatergic NMDAR activation using the glutamate site-specific antagonist D-(-)2-Amino-5-phosphonopentanoic acid (APV) (50 µM) (Fig. 1C-E). These findings were consistent in slices prepared from female mice (Fig. 1F-J). Due to off-target inhibition of AMPARs by 7-CK, we established that the amount of cDEP observed in the presence of the more selective and potent glycine site antagonist L689,560 (L689) (10 µM) was equivalent to its vehicle control (Fig. S1A-D) (Wong and Gray, 2018). Additionally, in agreement with other studies showing a role for NI-NMDAR signaling in LTD, 7-CK did not block the modest amount of LTD induced at naïve (unpotentiated) synapses (Fig. 2A-D) (Nabavi et al., 2013; Wong and Gray, 2018). Biological activity and sufficient concentrations of both 7-CK and L689 were demonstrated by confirming sLTP blockade (Fig. S2A-F). These pharmacological results are consistent with contrasting roles for ionotropic NMDAR signaling in sDEP and NI-NMDAR signaling in cDEP and LTD.

**Fig. 1.**
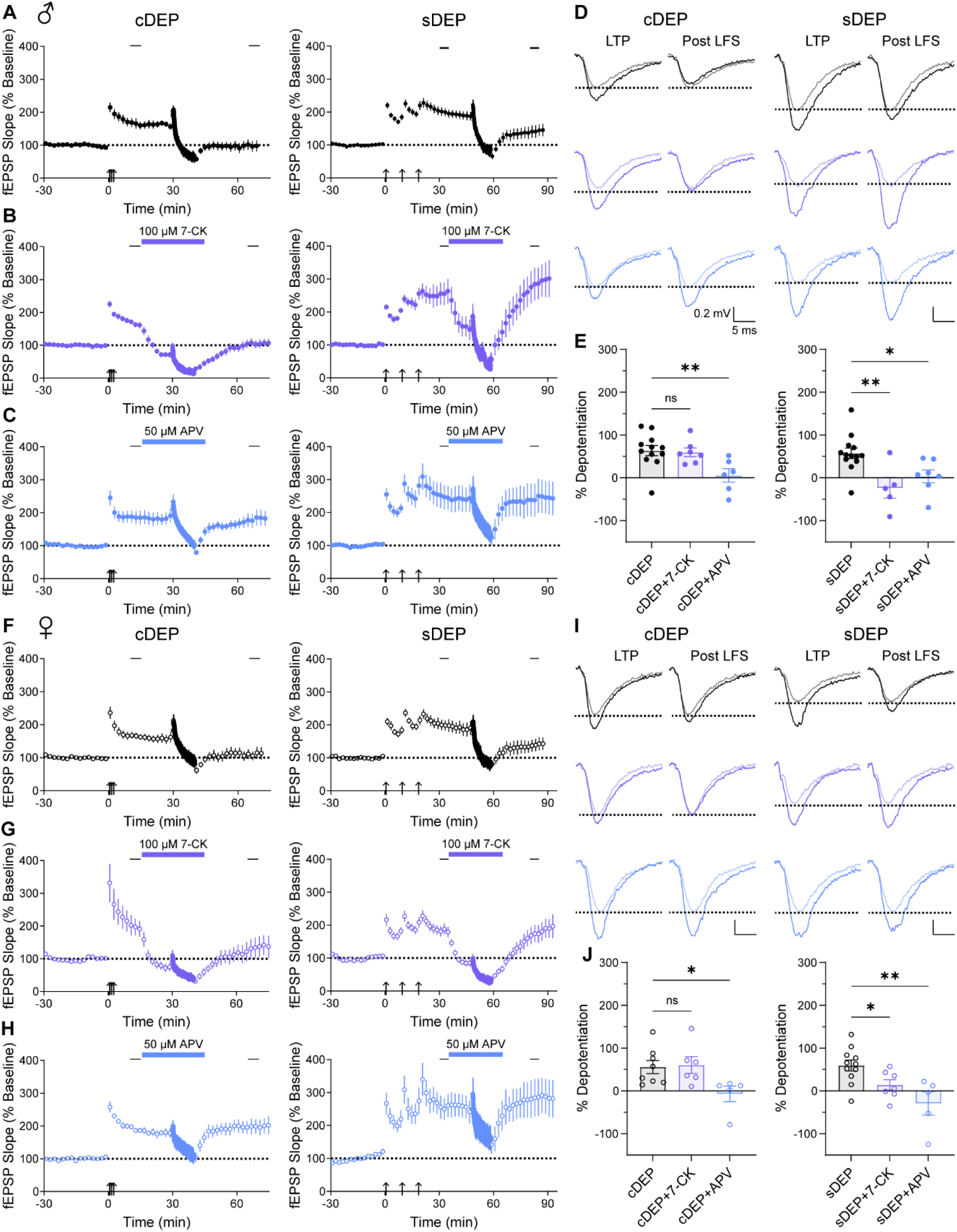
Ionotropic NMDAR signaling is required for sDEP but not cDEP in male and female mice. (**A**) fEPSPs slope as a percentage of baseline responses over time at Schaffer collateral-CA1 synapses with LTP induced using either a compressed or spaced induction paradigm followed by a 2Hz LFS 30 minutes later to initiate cDEP (left) or sDEP (right) in male mice (closed circles). Data are means ± SEM from 12 slices/11 mice (cDEP) and 13 slices/mice (sDEP). (**B**) sDEP, but not cDEP, was blocked by 100 µM 7-CK in male mice. Data are means ± SEM from 7 (cDEP) and 5 (sDEP) slices/mice. (**C**) 50 µM APV blocked both cDEP and sDEP in male mice. Data are means ± SEM from 6 slices/mice per group. (**D**) Individual representative field responses obtained at baseline, after LTP induction, and after LFS delivery for cDEP (left) and sDEP (right) in male mice in control conditions (black), in the presence of 7-CK (purple) or in the presence of APV (blue). Sample LTP and post LFS traces overlay a sample baseline trace with decreased opacity. Stimulus artifacts were excluded for clarity. (**E**) Average percent cDEP under control conditions versus in the presence of 7-CK or APV in male mice (left). Ordinary one-way ANOVA followed by Holm-Šídák post-hoc test, F = 5.42, p = 0.012; mean difference of cDEP vs. cDEP+7-CK = 4.0 ± 17.5%, p = 0.819; mean difference of cDEP vs. CDEP+APV = 58.1 ± 18.4% p = 0.0091. Average percent sDEP under control conditions, in the presence of 7-CK, or in the presence of APV in male mice (right). Ordinary one-way ANOVA followed by Holm-Šídák post-hoc test, F = 7.12, p = 0.0041; mean difference of sDEP vs. sDEP+7CK = 80.6 ± 23.5%, p = 0.0047; mean difference of sDEP vs. sDEP+APV = 53.5 ± 20.9%, p = 0.018. (**F**) A 2Hz LFS induced cDEP (left) and sDEP (right) in female mice (open circles). Data are means ± SEM from 8 slices/mice (cDEP) and 11 slices/10 mice (sDEP). (**G**) sDEP, but not cDEP, was blocked by 7-CK in female mice. Data are means ± SEM from 6 slices/5 mice (cDEP) and 7 slices/mice (sDEP). (**H**) APV blocked both cDEP and sDEP in female mice. Data are means ± SEM from 5 slices/mice per group. (**I**) Representative traces for cDEP (left) and sDEP (right) in female mice. (**J**) Average percent cDEP in female mice under control conditions versus in the presence of 7-CK or APV (left). Ordinary one-way ANOVA followed by Holm-Šídák post-hoc test, F = 4.03, p = 0.038; mean difference of cDEP vs. cDEP+7-CK = −4.6 ± 23.7%, p = 0.848; mean difference of cDEP vs. cDEP+APV = 62.8 ± 25.0%, p = 0.046. Average percent sDEP under control conditions versus in the presence of 7-CK or APV (right) in female mice. Ordinary one-way ANOVA followed by Holm-Šídák post-hoc test, F = 7.30, p = 0.0042; mean difference of sDEP vs. sDEP+7-CK = 45.3 ± 21.4%, p = 0.046; mean difference of sDEP vs. sDEP+APV = 88.5 ± 23.8%, p = 0.0027. Control, 7-CK and APV conditions are represented in black, purple and blue, respectively.

**Fig. 2.**
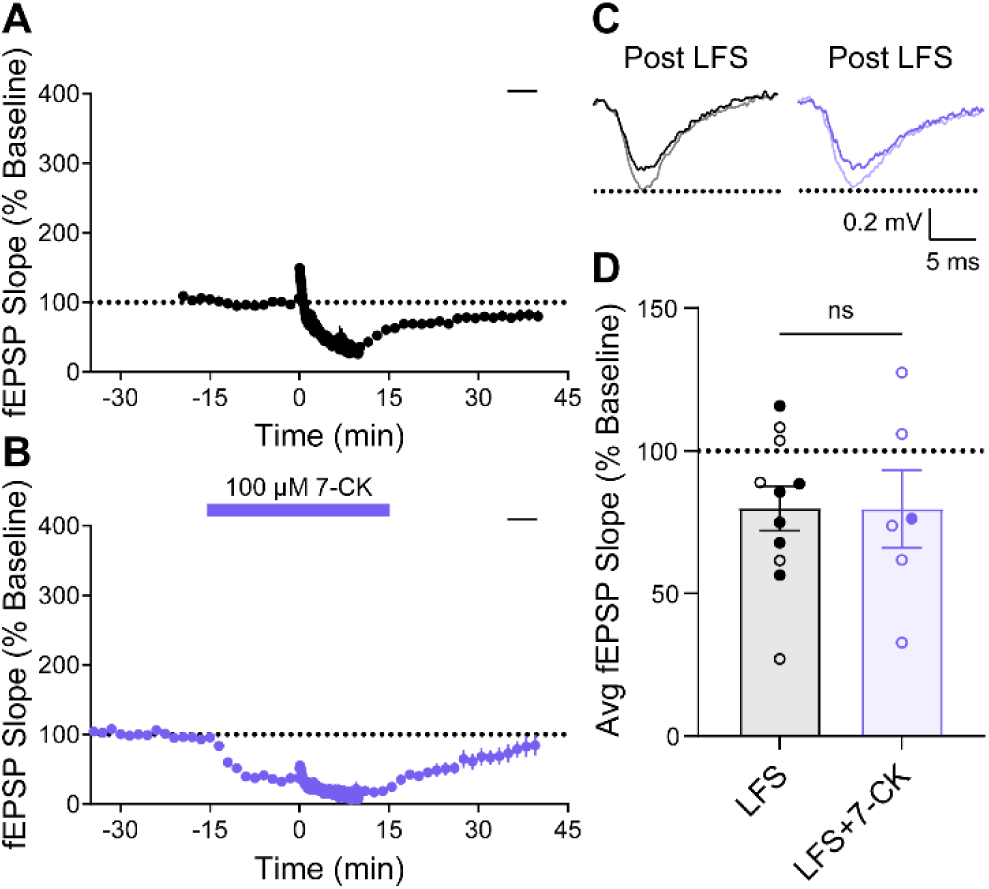
Long-term depression (LTD) does not require ionotropic NMDAR signaling. (**A**) At naïve synapses, LFS delivery led to a modest amount of LTD. Data are means ± SEM from 11 slices/mice. (**B**) LTD persisted in the presence of 100 µM 7-CK. Data are means ± SEM from 6 slices/mice. (**C**) Representative field responses obtained at baseline and after LFS delivery in the absence (black) or presence (purple) of 7-CK. (**D**) Average percent LTD under control conditions versus in the presence of 7-CK obtained at the end of recording, as indicated by the black bars in (A) and (B) (unpaired Student’s t test, mean difference = 0.2 ± 14.5%, p = 0.990). Data from male (closed circles) and female (open circles) mice combined.

### NI-NMDAR signaling is required for cDEP only

We have recently developed a cell permeable peptide designed to disrupt NI-NMDAR signaling (Zhang et al., 2023). The transactivator of transcription (Tat)-conjugated peptide (C1.1) mimics a region of the C1 domain in the GluN1 C-terminal to interfere with protein interactions necessary for the intracellular propagation of NI-NMDAR signaling (Warnet et al., 2020). Similar mimetic peptides have also been employed by other groups to disrupt protein interactions with the NMDAR C-terminal, presumably by competing for binding targets (Cahill et al., 2014; Sheehan et al., 2024). We empirically determined a peptide concentration (50 uM) that did not affect postsynaptic responses (Fig. 3A) and bath-applied this concentration of peptide to assess the necessity of NI-NMDA signaling in DEP. As expected, the scrambled control peptide, C1.1Scr, did not prevent cDEP or sDEP (Fig. 3B). However, we observed that on average, inhibiting NI-NMDAR signaling with C1.1 completely blocked cDEP (Fig. 3C left, D, E). In contrast, sDEP persisted in the presence of C1.1 (Fig. 3C right, D, E). These results confirm a requirement for NI-NMDAR signaling in cDEP only.

**Fig. 3.**
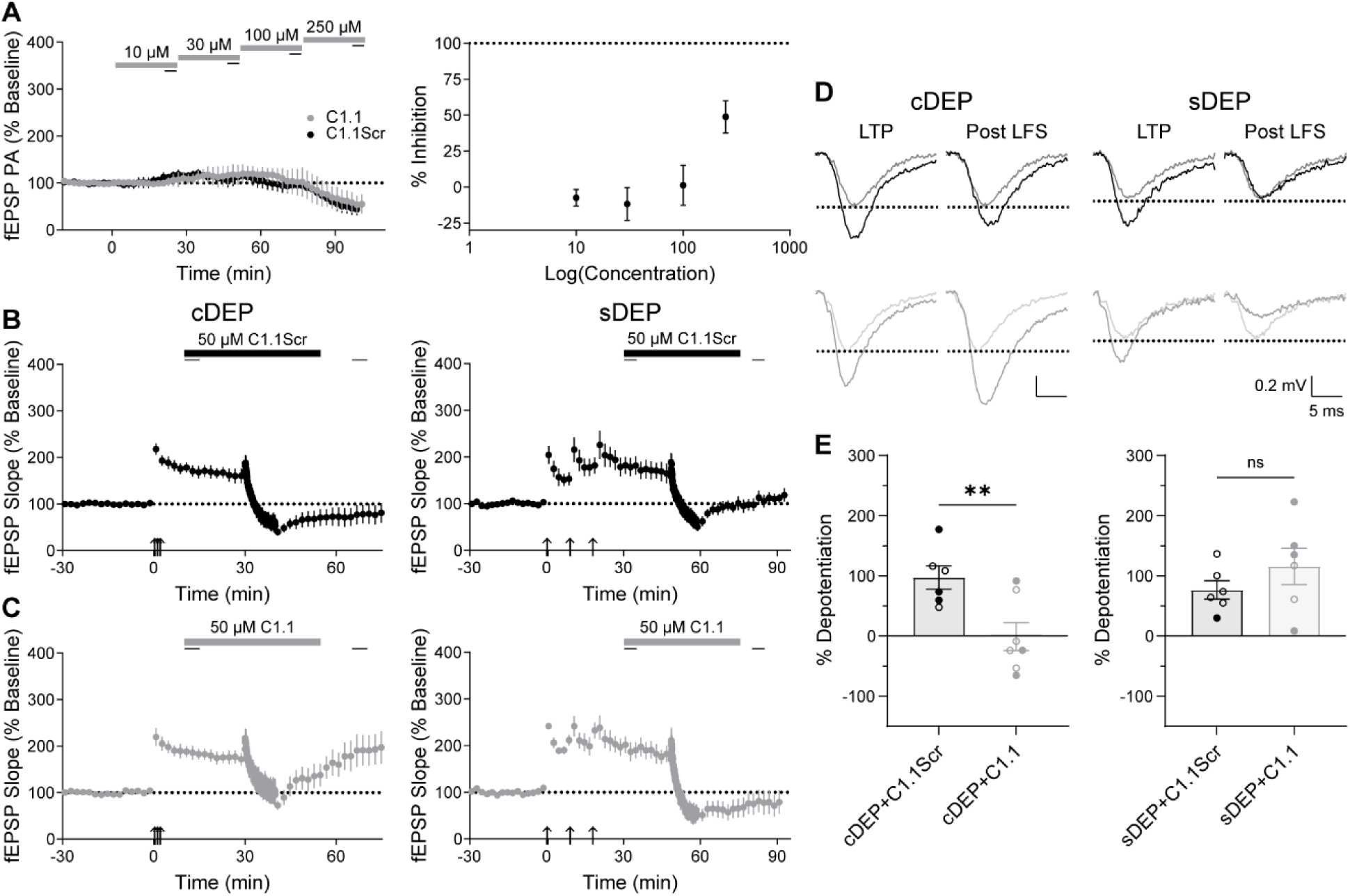
Preventing NI-NMDAR signaling with the cell-permeable C1.1 peptide blocks cDEP but not sDEP. (**A**) In the absence of plasticity, both C1.1 and its scrambled control (C1.1Scr) dampened synaptic responses at high concentrations (>100 µM) as shown by reductions in the fEPSP peak amplitude (PA) with progressive peptide addition over time (left). A corresponding logarithmic concentration curve is shown on the right, where the percentage inhibition was calculated using 5-minute averages indicated by the black bars. Data are means ± SEM from 3 slices/mice per group. (**B**) Both cDEP (left) and sDEP (right) persisted in the presence of 50 µM of the scrambled control peptide, C1.1Scr. Data are means ± SEM from 6 slices/mice per group. (**C**) 50 µM of C1.1 applied during LFS administration blocked cDEP (left) but not sDEP (right). Data are means ± SEM from 7 (cDEP) and 6 (sDEP) slices/mice per group. (**D**) Representative field responses obtained at baseline, after LTP induction, and after LFS delivery for cDEP (left) and sDEP (right) in the presence of C1.1Scr (black) or C1.1 (grey). (**E**) Average percent cDEP in the presence of C1.1Scr versus C1.1 (unpaired Student’s t test, mean difference = 98.2 ± 30.9, p = 0.0088) (left). Average percent sDEP in the presence of C1.1Scr versus C1.1 (unpaired Student’s t test, mean difference = −39.2 ± 33.9%, p = 0.275) (right). Data from male (closed circles) and female (open circles) mice combined.

To assess possible presynaptic contributions during DEP, we monitored the percent change in paired-pulse ratios (PPRs) of fEPSPs evoked 50 ms apart after DEP compared to LTP, normalized to the baseline PPR. No statistically significant differences were observed in the percent change in PPR in the presence versus absence of APV, or C1.1 versus C1.1Scr (Fig. S3A-D). These observations were consistent following both cDEP and sDEP and are congruent with a postsynaptic locus of plasticity expression. However, further research is required to elucidate presynaptic alterations that may occur during LTP and DEP.

### GluA1/GluA2 AMPAR subunit ratio is enhanced following cLTP compared to sLTP

So far, we have shown that ionotropic and non-ionotropic NMDAR signaling pathways are uniquely recruited during DEP of sLTP and cLTP. Downstream of NMDARs, AMPAR internalization is necessary for hippocampal DEP (Migues et al., 2016) and AMPAR expression is differentially modulated during LTD and DEP (Zhu et al., 2005). The hippocampus consists primarily of GluA1/GluA2 AMPARs, with a small proportion of GluA2-lacking, Ca^2+^-permeable AMPARs that contribute to certain types of plasticity depending on the method of induction (Park et al., 2016; Park et al., 2018). Therefore, we probed for alterations in the GluA1/GluA2 ratio following plasticity induction. Using permeabilized immunostaining on slices fixed in 4% paraformaldehyde after electrophysiology recordings, we quantified the amount of GluA1 and GluA2 based on the fluorescence intensity in the stratum radiatum (SR). After normalization to a naïve slice stained and imaged at the same time, we found a small but statistically significant increase in the ratio of GluA1/GluA2 fluorescence intensity following cLTP compared to sLTP (Fig. 4A, B). In contrast, the average GluA1/GluA2 fluorescence ratio was equivalent following cDEP compared to sDEP (Fig. 4C). To assess the spatial distribution of GluA1 and GluA2, we generated intensity profiles along the normalized length of the SR (Fig. 4D-F). A statistically significant main effect of LTP type was observed when comparing the fluorescence intensity of GluA1 along the SR, relative to the maximal intensity within each slice (Fig. 4E). No detectable differences were found in the normalized GluA1 or GluA2 intensity profiles following cDEP compared to sDEP (Fig. 4F). These results imply that the distribution of GluA1 may differ along the length of the SR following sLTP compared to cLTP. We conclude that although AMPAR subunit expression may be differentially altered following cLTP and sLTP induction, cDEP and sDEP appear to converge at the level of AMPAR expression.

**Fig. 4.**
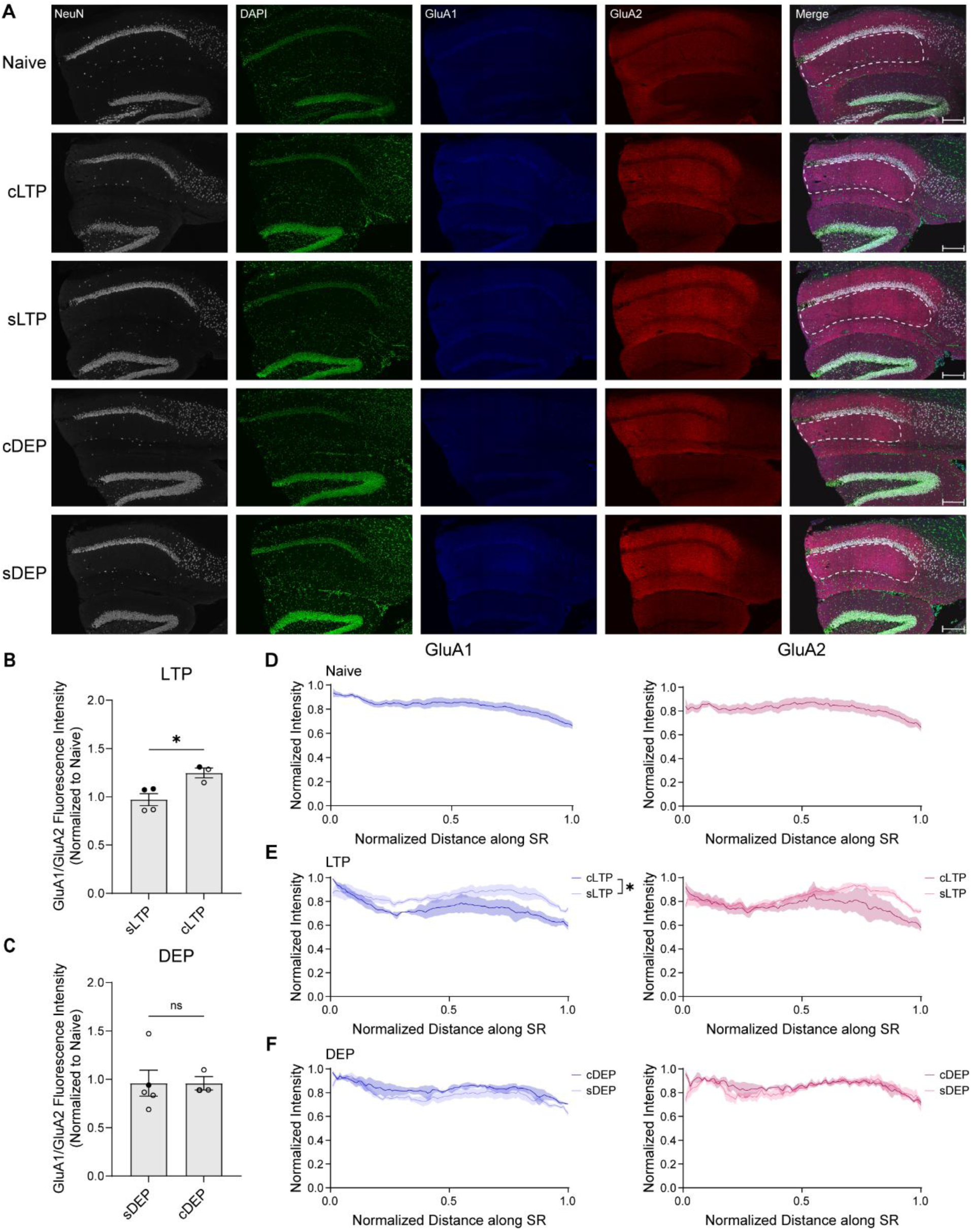
cLTP and sLTP are associated with different AMPAR subunit expression. (**A**) Representative confocal microscopy images of hippocampal slices following electrophysiology stained for NeuN (grey, AF-647), GluA1 (blue, AF-488), GluA2 (red, AF-594), and DAPI (green) and corresponding merged images for naïve, cLTP, sLTP, cDEP, and sDEP slices (maximum intensity projection). The dotted white line shown in the merged image represents the region of interest (stratum radiatum, SR) for each slice. Scalebar = 200 µm. (**B**) After normalization to naïve slices, the GluA1/GluA2 ratio in the SR was higher following cLTP compared to sLTP (unpaired Student’s t test, mean difference = 0.277 ± 0.084 times the naïve GluA1/GluA2 ratio, p = 0.022). Data are means ± SEM from 4 (sLTP) and 3 (cLTP) biological replicates. (**C**) No differences in the GluA1/GluA2 ratio were observed following cDEP compared to sDEP (unpaired Student’s t test, mean difference = −0.002 ± 0.186 times the naïve GluA1/GluA2 ratio, p = 0.993). Data are means ± SEM from 5 (sDEP) and 3 (cDEP) biological replicates. (**D**) Fluorescence intensity profiles normalized to the maximum fluorescence intensity as a function of distance along the SR (normalized to 1) for GluA1 (left) and GluA2 (right) in naïve slices. Data are means ± SEM from 10 biological replicates. (**E**) Normalized fluorescence intensity profiles for GluA1 (left) and GluA2 (right) in slices fixed after cLTP or sLTP. Data are means ± SEM from 4 (sLTP) and 3 (cLTP) biological replicates. Differences in GluA1 fluorescence intensity along the SR relative to the maxima were observed between cLTP and sLTP (mixed effects model, distance x LTP F(99, 494) = 1.11, p = 0.239; main effect of LTP type F(1,5) = 9.43, p = 0.028). No statistically significant differences in GluA2 distribution were observed between cLTP and sLTP (mixed effects model, distance x LTP F(99, 494) = 1.23, p = 0.081, LTP type F(1, 5) = 3.11, p = 0.138). (**F**) Normalized fluorescence intensity profiles for GluA1 (left) and GluA2 (right) in slices fixed after cDEP or sDEP. Data are means ± SEM from 5 (sDEP) and 3 (cDEP) biological replicates. No statistically significant differences were observed between cDEP and sDEP for GluA1 (distance x DEP F(99, 591) = 0.459, p > 0.999, DEP type F(1, 6) = 2.07, p = 0.200) or GluA2 (distance x DEP F(99, 591) = 0.680, p = 0.991, DEP type F(1, 6) = 1.69, p = 0.242) using mixed effects analysis. Data from male (closed circles) and female (open circles) mice combined, 1-2 technical replicates per biological replicate.

### sDEP and cDEP are associated with distinct GluA1 phosphorylation states

Phosphorylation of the GluA1 subunit of the AMPAR can affect channel properties, kinetics and localization at the synapse. Different phosphatases are recruited during synaptic weakening depending on the presence of prior LTP induction (Zhuo et al., 1999; Jouvenceau et al., 2003). Further, distinct AMPAR phosphorylation states are associated with LTD and DEP, suggesting that LTP induction may have a metaplastic effect on GluA1 phosphorylation (Lee et al., 2000). Dephosphorylation of GluA1 at serine S831 and S845 have been linked with plastic modifications by controlling single channel AMPAR conductance (Derkach et al., 1999) and open channel probability/AMPAR endocytosis, respectively (Banke et al., 2000; Man et al., 2007). To test whether unique AMPAR phosphorylation states may accompany sDEP and cDEP, we performed SDS-PAGE and immunoblotting on hippocampal slices flash frozen 30-40 minutes after electrophysiological DEP induction alone or in the presence of 7-CK or APV. After isolating the hippocampus, a whole lysate preparation was used to probe phosphorylated GluA1 at sites S831 and S845 as well as total GluA1 and GluA2 in individual slices. No differences were observed in the total amount of GluA1 or GluA2 following cDEP compared to sDEP induced in the absence or presence of 7-CK/APV after normalizing to GAPDH within lanes and to naïve slices within blots (Fig. 5A, B). Notably, the normalized amount of phosphorylated GluA1 (pGluA1) at S831 was elevated following cDEP compared to sDEP in the absence or presence of 7-CK (Fig. 5C). Similarly, pGluA1 at S845 was lower following sDEP compared to cDEP in control conditions (Fig. 5D). Within DEP conditions, pGluA1 S845 was elevated following sDEP induction in the presence of 7-CK relative to control conditions. In contrast, application of C1.1 compared to C1.1Scr during cDEP did not result in any statistically significant differences in GluA1, GluA2 or pGluA1 at S831 or S845 (Fig. 5E-H; see Fig. S4A-C for full blots). Together, our observations suggest that GluA1 phosphorylation may be differentially promoted following cDEP compared to sDEP.

**Fig. 5.**
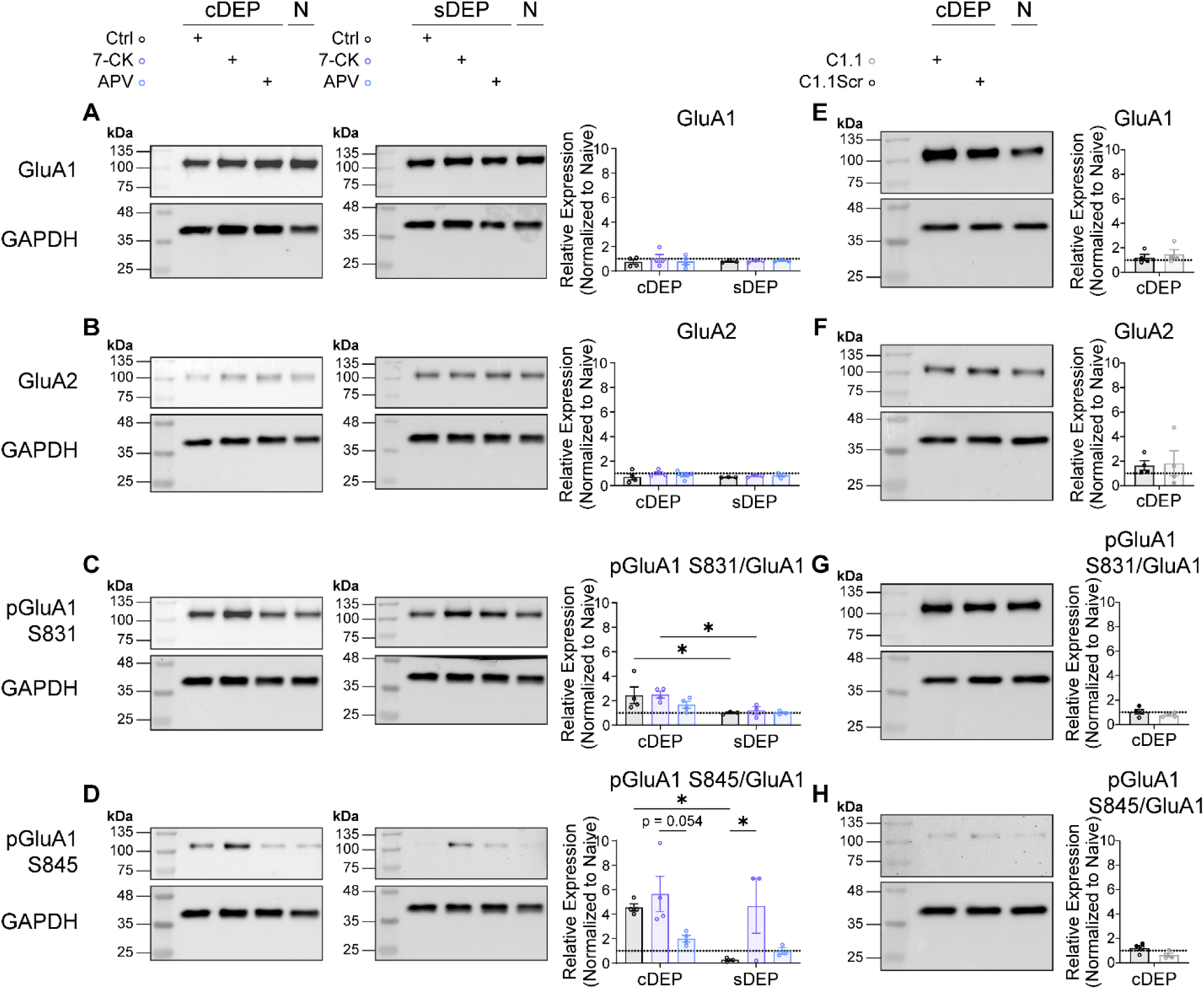
GluA1 phosphorylation is differentially regulated following cDEP and sDEP. (**A**) GluA1 normalized to naïve hippocampal slices following cDEP and sDEP in control conditions (black), in the presence of 100 µM 7-CK (purple) and in the presence of 50 µM APV (blue). DEP x treatment F(2, 15) = 0.240, p = 0.790. (**B**) GluA2 normalized to naïve slices following cDEP and sDEP in control conditions and in the presence of 7-CK or APV. DEP x treatment F(2, 15) = 0.220, p = 0.805. (**C**) pGluA1 S831/GluA1 ratio normalized to naïve hippocampal slices following cDEP and sDEP in control conditions and in the presence of 7-CK or APV. DEP x treatment F(2, 15) = 0.583, p = 0.571, main effect of DEP type F(1, 15) = 12.27, p = 0.003. Post-hoc cDEP+7-CK vs. sDEP+7-CK least squares (LS) mean difference = 1.3 ± 0.6 times naïve pGluA1 S831/GluA1 expression, p = 0.034; cDEP vs. sDEP LS mean difference = 1.4 ± 0.6 times naïve pGluA1 S831/GluA1 expression, p = 0.021. (**D**) pGluA1 S845/GluA1 ratio normalized to naïve hippocampal slices following cDEP and sDEP in control conditions and in the presence of 7-CK or APV. DEP x treatment F(2, 15) = 1.62, p = 0.231, main effect of DEP type (F(1, 15) = 5.88, p = 0.028) and drug treatment (F(2, 15) = 6.57, p = 0.0089). Post-hoc cDEP+7CK vs. cDEP+APV LS mean difference = 3.7 ± 1.4 times naïve pGluA1 S845/GluA1 expression, p = 0.054; sDEP vs. sDEP+7-CK LS mean difference = −4.4 ± 1.6 times naïve pGluA1 S845/GluA1 expression, p = 0.043; cDEP vs. sDEP LS mean difference = 4.3 ± 1.5 times naïve pGluA1 S845/GluA1 expression, p = 0.012. Data in (A-D) are means ± SEM from 4 (cDEP) and 3 (sDEP) biological replicates. (**E-H**) AMPAR expression and phosphorylation states are unaltered following cDEP in the presence of C1.1 compared to C1.1Scr. (**E**) GluA1 normalized to naïve hippocampal slices following cDEP in the presence of C1.1Scr (black) versus C1.1 (grey) (p = 0.555). (**F**) GluA2 normalized to naïve hippocampal slices following cDEP in the presence of C1.1Scr versus C1.1 (p = 0.876). (**G**) pGluA1 S831/GluA1 ratio normalized to naïve hippocampal slices following cDEP with C1.1Scr versus C1.1 (p = 0.238) (**H**) pGluA1 S845/GluA1 ratio normalized to naïve hippocampal slices following cDEP with C1.1Scr versus C1.1 (p = 0.067). Data in (E-G) are means ± SEM from 4 biological replicates per group. 1-2 technical replicates per biological replicate. Blots were cut to probe for each protein (see Fig. S5A-C). Each drug treatment condition was normalized to a naïve slice run in the same blot. Statistical comparisons were made using ordinary two-way ANOVA followed by Holm-Šídák post-hoc comparisons within and across drug treatment conditions (A-D) or unpaired Student’s t test (E-H) as appropriate.

### Dendritic Arc mRNA trafficking is uniquely required during sDEP

Due to the differences we observed in AMPAR expression and phosphorylation, we explored the mechanisms that may mediate AMPAR endocytosis during sDEP and cDEP. The activity-regulated cytoskeleton-associated protein (Arc/Arg3.1) has been identified as a key mediator of synaptic plasticity (Zhang and Bramham, 2021). Arc is trafficked to dendrites following neuronal activity and mobilizes AMPARs to and from the synapse (Chowdhury et al., 2006; Carmichael and Henley, 2018). Recently, it has been revealed that Arc couples to NMDARs to constrain DEP but not LTD (Yang et al., 2023). Using an *Arc^CreER^*mouse model in which dendritic Arc mRNA trafficking is specifically disrupted by a Cre recombinase insertion (CreER^T2^) at the 3’ untranslated region (UTR) of the *Arc* loci (JAX:021881) (Kobayashi et al., 2005), we tested the requirement for Arc in cDEP and sDEP. *Arc^CreER^*mice exhibited alterations in baseline pre- and postsynaptic functioning compared to WT mice (Fig. 6A). *Arc^CreER^* mice also exhibited impaired dendritic Arc mRNA trafficking, which we confirmed using reverse transcription polymerase chain reaction (RT-qPCR) of RNA isolated from the whole hippocampus of homozygous *Arc^CreER^* and wildtype (WT) mice (Fig. 6B). Despite these alterations, short-term sLTP and cLTP remained intact in *Arc^CreER^* slices (Fig. 6C, D), consistent with other genetic models in which Arc is disrupted (Kyrke-Smith et al., 2021; Yang et al., 2023). Importantly, sDEP, but not cDEP, was markedly impaired in *Arc^CreER^*mice (Fig. 6C-F). We interpret these findings as evidence that in addition to recruiting distinct forms of NMDAR signaling and AMPAR phosphorylation states, cDEP and sDEP may also have disparate requirements for Arc-mediated synaptic AMPAR removal.

**Fig. 6.**
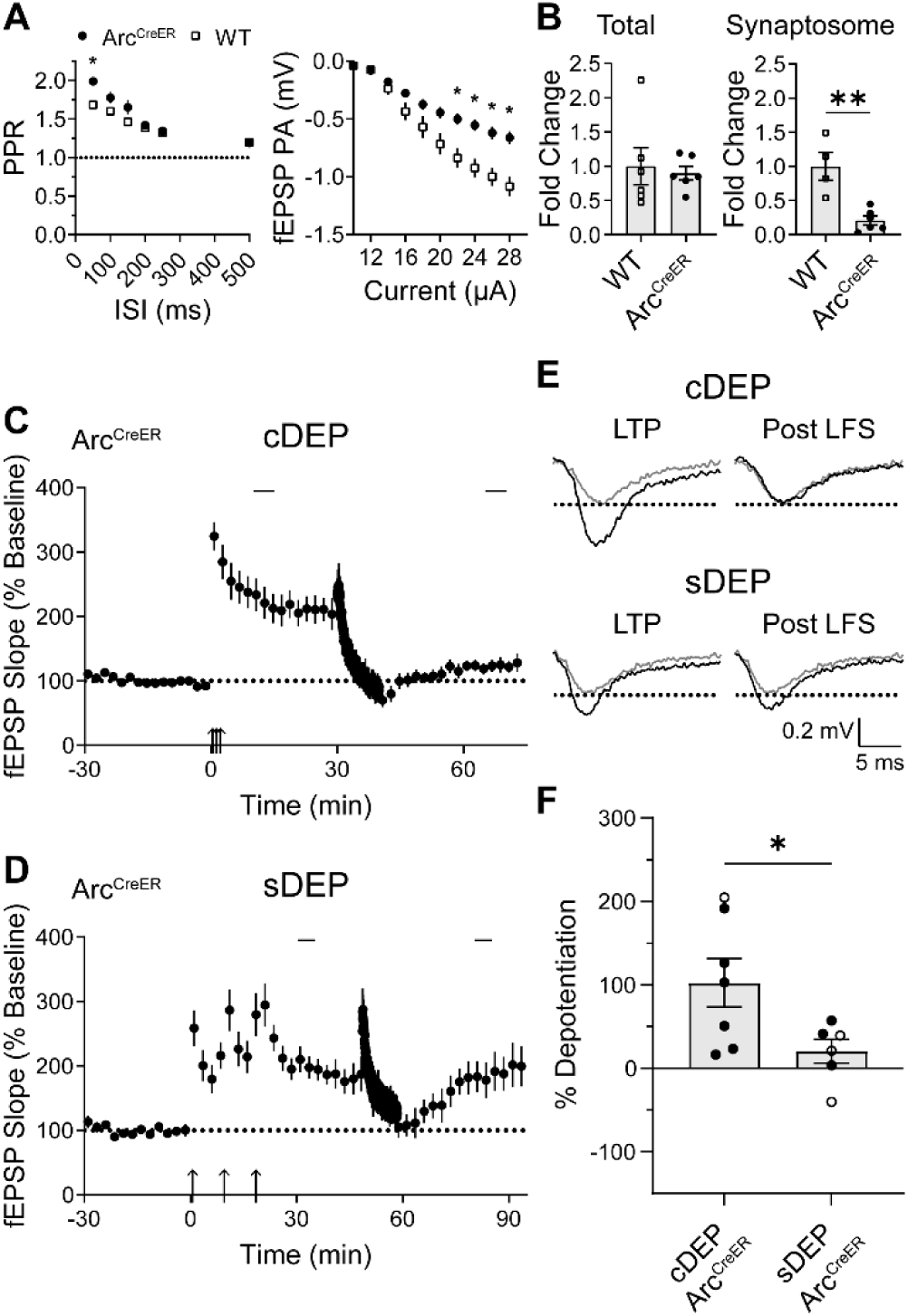
sDEP, but not cDEP, is impaired in *Arc^CreER^* mice. (**A**) In naïve hippocampal slices, the paired pulse ratio (PPR) was elevated in *Arc^CreER^* mice (circles, n = 4 slices/mice) compared to WT mice (squares, n = 5 slices/mice) (repeated measures two-way ANOVA followed by Holm-Šídák post-hoc comparisons between genotypes, ISI x genotype F(5, 35) = 7.37, p < 0.0001; *Arc^CreER^* vs. WT at 50 ms ISI p = 0.014) (left). The peak amplitude (PA) as a function of stimulus intensity (input/output curve) was reduced in *Arc^CreER^* mice (n = 8 slices/mice) compared to WT mice (n = 8 slices/mice) (repeated measures two-way ANOVA followed by Holm-Šídák post-hoc comparisons between genotypes, current x genotype F(9, 126) = 13.62, p<0.0001; *p<0.05 as indicated for post-hoc between genotype comparisons) (right). (**B**) WT and *Arc^CreER^* mice had equivalent levels of total Arc mRNA, quantified using RT-qPCR on isolated hippocampus samples (unpaired Student’s t test, mean difference = 0.10 ± 0.29 p = 0.732) (left). However, *Arc^CreER^* mice exhibited reduced synaptosomal Arc mRNA levels compared to WT mice (unpaired Student’s t test, mean difference = −0.79 ± 0.18, p = 0.0024) (right). Data are means ± SEM from 6 male mice per group (total Arc mRNA), and 4 and 6 male mice (synaptosomal Arc mRNA). (**C**) cLTP was readily induced and depotentiated in *Arc^CreER^* mice. Data are means ± SEM from 7 slices/5 mice. (**D**) sLTP was readily induced but sDEP was impaired in *Arc^CreER^* mice, on average. Data are means ± SEM from 6 slices/mice. (**E**) Representative field responses obtained at baseline, after LTP, and after LFS delivery for cDEP (top) and sDEP (bottom). (**F**) Average percent DEP of cLTP versus sLTP in *Arc^CreER^* mice (unpaired Student’s t test, mean difference = 82.0 ± 34.1%, p = 0.035). Data from male (closed circles) and female (open circles) combined.

## Discussion

We examined whether the temporal spacing of LTP induction affects the mechanisms invoked during NMDAR-mediated synaptic DEP in the hippocampus. Our results reveal distinct ionotropic and non-ionotropic NMDAR signaling pathways recruited during DEP depending on the type of LTP induced. Specifically, we observed that non-ionotropic NMDAR signaling is uniquely required for DEP of cLTP in both male and female mice. In contrast, ionotropic NMDAR signaling is necessary to induce DEP after the induction of sLTP. Downstream of NMDAR signaling, sDEP and cDEP converge at the level of AMPAR expression. However, the mechanisms by which synaptic strength is reduced following DEP may differ, as sDEP uniquely requires Arc and appears to exhibit reduced GluA1 phosphorylation relative to cDEP. The results of this study are consistent with an interpretation of metaplastic tags that are set during plastic events underlying learning and which influence the susceptibility of a memory trace to disruption and forgetting.

Our main findings corroborate previous reports of NI-NMDAR signaling in LTD and DEP (Nabavi et al., 2013; Latif-Hernandez et al., 2016). However, we found that the type of LTP induction governed the requirement for ionotropic or non-ionotropic NMDAR signaling during DEP. Leveraging a cell-permeable peptide, we disrupted effector interactions with the C tail of the NMDAR necessary for NI-NMDAR signaling (Zhang et al., 2023), allowing us to probe the necessity of NI-NMDAR signaling and reveal that NI-NMDAR signaling is involved in cDEP, but not sDEP. The precise NMDAR binding partner interactions disrupted by C1.1 are not yet established. Possible interactions may include PP1 tethered by the A-kinase anchoring protein Yotiao, known to be involved in NI-NMDAR signaling (Westphal et al., 1999; Aow et al., 2015), or G-protein-coupled receptors such as the dopamine D1 receptor (Cahill et al., 2014).

Postsynaptic Ca^2+^ influx can activate a multitude of intracellular signaling cascades to tune stimulus responses through structural and functional modifications. Our results suggest that NMDAR-mediated Ca^2+^ influx may be required to overcome a heightened plasticity threshold set during sLTP induction. The same LFS is used to induce cDEP and sDEP suggesting that the degree of Ca^2+^ influx through the NMDAR and the downstream pathways recruited during DEP may vary depending on the molecular landscape of the synapse. For example, postsynaptic NMDAR composition can be modified by synaptic activity and alters subsequent plasticity induction thresholds (Xu et al., 2009; Foster et al., 2010; Ge et al., 2019). Our finding that NI-NMDAR signaling is involved in both cDEP and LTD at naïve synapses suggests that synapses may be primed for ionotropic NMDAR signaling following sLTP induction. The precise mechanisms by which sLTP induction confers a bias towards ionotropic NMDAR signaling, and the downstream effectors necessary for sDEP and cDEP remain to be determined.

AMPAR expression is dynamically regulated to amplify or dampen postsynaptic responses following plasticity induction. Although we did not find differences in the GluA1/GluA2 ratio in the SR following cDEP and sDEP, we observed an increase in GluA1/GluA2 following cLTP compared to sLTP. Transient insertion of Ca^2+^-permeable AMPARs at perisynaptic sites is thought to be triggered upon delivery of a TBS (Park et al., 2016). During sLTP, Ca^2+^-permeable AMPARs are replaced by more stable GluA2-containing AMPARs upon subsequent spaced TBS events (Park et al., 2016; Park et al., 2021). Therefore, our observation of an elevated GluA1/GluA2 ratio in the SR following cLTP may have been a consequence of perisynaptic priming of GluA2-lacking, Ca^2+^-permeable AMPARs. In contrast, unstable GluA2-lacking AMPARs may have been replaced with more stable GluA2-containing receptors following sLTP induction at the time of fixation. Further, our findings hint at possible differences in the spatial distribution of AMPAR subunits following cLTP and sLTP along the SR, however the reasons for these differences remain to be investigated. After cDEP or sDEP induction, no differences in AMPAR expression were observed. As such, we conclude that DEP converges at the level of AMPAR subunit expression.

We observed comparable total hippocampal GluA1 and GluA2 expression following sDEP and cDEP in both the presence and absence of NMDAR antagonists that block DEP. However, pGluA1 at both S831 and S845 was elevated following cDEP compared to sDEP. pGluA1 S845 was also enhanced in the presence of 7-CK compared to control conditions during sDEP, consistent with sDEP blockade via 7-CK. In the presence of 7-CK, pGluA1 phosphorylation at S831 was higher after cDEP compared to sDEP, consistent with opposing effects of 7-CK on the two types of DEP. As expected, no differences in GluA1 phosphorylation were observed between sDEP and cDEP in the presence of APV, which functionally blocks both types of DEP. AMPAR subunit expression and phosphorylation levels were indistinguishable following cDEP induced in the presence of C1.1 compared to its scrambled control. It is possible that although C1.1 blocked the expression of DEP, this peptide may not prevent all downstream effects of DEP induction. Additionally, the differences observed between sDEP and cDEP may have arisen from either enhanced dephosphorylation of GluA1 during sDEP or opposing GluA1 phosphorylation and dephosphorylation during cDEP and sDEP, respectively. Others have previously reported GluA1 dephosphorylation at the CaMKII/PKC binding site (S831) during DEP (Lee et al., 2000; Huang et al., 2001). AMPAR dephosphorylation at the GluA1 PKA binding site (S845) has instead been observed following LTD (Lee et al., 2000). However, certain types of LTD-inducing stimuli have also been shown to increase phosphorylation of GluA1 at both S831 and S845 (Delgado and O’dell, 2005), or simultaneously increase S831 and decrease S845 phosphorylation (Ai et al., 2011). The resulting effects of S831/S845 phosphorylation or dephosphorylation during sDEP and cDEP, and the relationship with NMDAR signaling pathways are intriguing questions for further study. It should be noted that 7-CK application during DEP may have had off-target effects on GluA1 phosphorylation, as it is known to partially inhibit AMPARs as well as NMDARs (Wong and Gray, 2018). Overall, our immunoblotting results suggest that phosphatase recruitment or activity may be differentially recruited following cDEP and sDEP.

Finally, sDEP was diminished in an *Arc^CreER^* mouse model with impaired dendritic Arc mRNA trafficking. Conversely, cDEP was depotentiated to baseline levels in *Arc^CreER^* slices, as previously demonstrated by others in a constitutive Arc knockout model (Yang et al., 2023). These results suggest that the different pathways involved in sDEP and cDEP may modulate synaptic plasticity by regulating AMPAR kinetics and/or endocytosis. However, we cannot rule out the possibility that different mechanisms are recruited during LTP and/or DEP in *Arc^CreER^* mice due to altered pre- and postsynaptic functioning compared to WT mice. Nonetheless, the metaplastic regulation of Arc may explain its ability to promote or restrict synaptic weakening under different conditions (Wall et al., 2018; Yang et al., 2023). Indeed, our findings are consistent with previous reports of an elevated proportion of CA1 neurons expressing Arc mRNA following spaced compared to massed exposure to a novel environment (Guzowski et al., 2006).

We propose a model in which sDEP requires ionotropic NMDAR signaling and downstream trafficking of Arc to activated synapses for AMPAR endocytosis. Conversely, NI- NMDAR signaling is involved in the induction of cDEP where plasticity expression is independent of Arc-mediated AMPAR endocytosis. Accordingly, NMDAR activation and Ca^2+^ influx during plasticity events has been shown to induce Arc translation and transcription (Bloomer et al., 2008; Zheng et al., 2009; Carmichael and Henley, 2018) (however see (Chen et al., 2017)) and Arc-NMDAR association (Yang et al., 2023). During sDEP, NMDAR-mediated Ca^2+^ influx may provide an additional plasticity signal by activating Ca^2+^-dependent effectors including phosphatases that dephosphorylate AMPARs to reduce open channel probability and/or promote endocytosis via Arc. During cDEP, NI-NMDAR signaling likely recruits distinct molecular players that act to reduce synaptic transmission in an Arc-independent manner. The differential recruitment of NMDAR pathways following cLTP and sLTP induction we have shown may also aid in reconciling contradictory evidence for ionotropic or non-ionotropic NMDAR signaling in LTD, which could depend on the history of synaptic activity or other experimental conditions.

The metaplastic requirement for ionotropic/non-ionotropic NMDAR signaling and Arc during DEP of sLTP and cLTP may extend to the physiological regulation of forgetting. The parameters by which LTP is induced may define a metaplastic role that promotes the preservation of synaptic plasticity and encoding of certain memories while targeting others for degradation (Hardt et al., 2014; Zhang et al., 2018). Consistent with our in vitro findings, temporally spaced behavioural training results in more robust memories and evokes different forgetting mechanisms than massed training paradigms (Bello-Medina et al., 2013; Jiang et al., 2016). Pharmacologically promoting ionotropic NMDAR signaling may enable the disruption of memories ordinarily resistant to forgetting in pathological states like posttraumatic stress disorder. Conversely, selectively targeting NI-NMDAR signaling may result in the preservation of weak memories during pathological forgetting. In fact, NI-NMDAR signaling has already been associated with βamyloid-induced synaptic depression (Kessels et al., 2013) and excitotoxic cell death (Weilinger et al., 2016). Together, our results highlight the ability of the synapse to ‘remember’ prior activity which uniquely influences the mechanisms recruited during subsequent plasticity events.

## Acknowledgments

We would like to thank Dr. Patrick Tidball (University of Toronto, Toronto, Canada) for electrophysiology guidance, Dr. Zhengping Jia (The Hospital for Sick Children, Toronto, Canada) for assistance with developing an immunoblotting protocol, and Dr. Michael E. Hildebrand (Carleton University, Ottawa, Canada) for consultation on analysis.

## Author contributions

Conceptualization: QP, AJR, RPB

Formal analysis: QP, JA, SWF

Investigation: QP, JA, SWF

Resources: RPB

Writing – original draft: QP

Writing – review & editing: QP, JA, SWF, AJR, RPB

Visualization: QP

Supervision: RPB

Funding acquisition: RPB

## Data and materials availability

All data are available in the main text or the supplementary materials. Original data are available upon reasonable request from the authors.

## Supplementary Materials

### Supplementary Figures

**Fig. S1.**
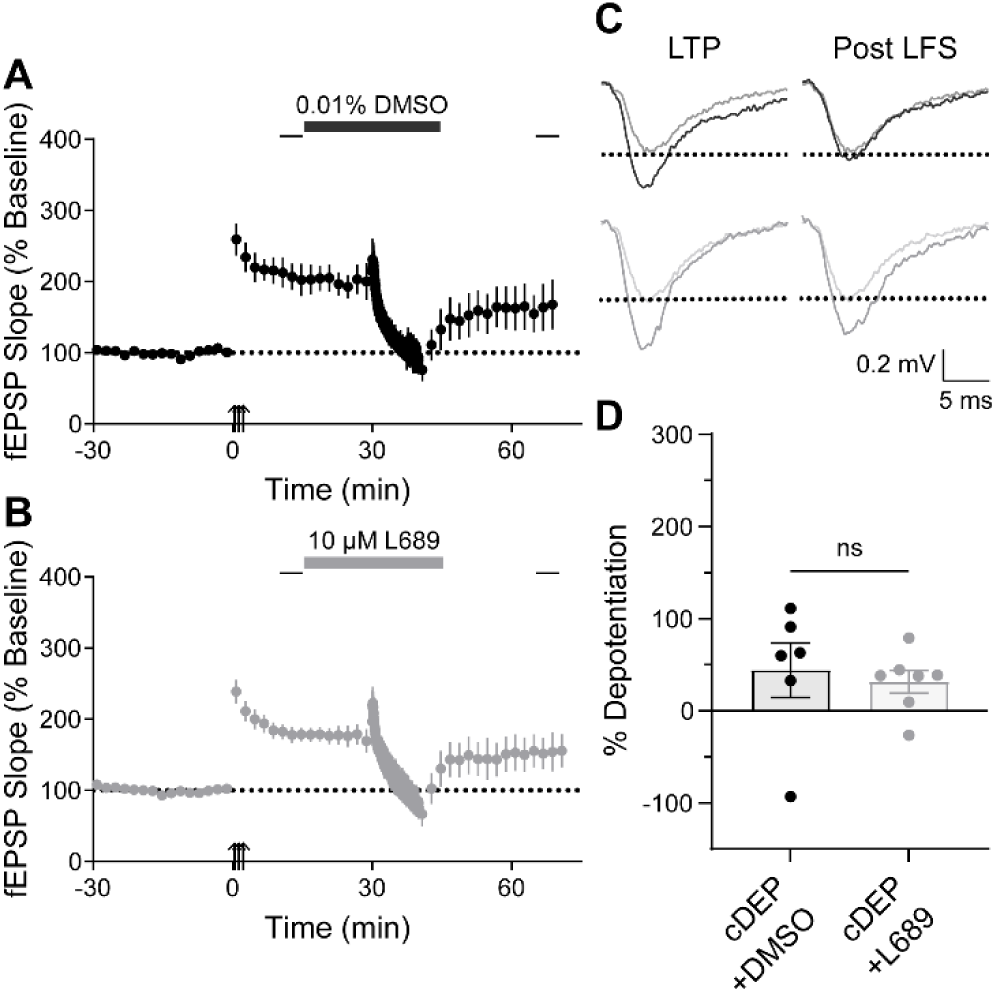
cDEP is not blocked by the NMDAR glycine site antagonist L689. (**A**) Following cLTP induction, a 2Hz LFS induced DEP in the presence of vehicle alone (0.01% DMSO) or (**B**) 10 µM L689 dissolved in DMSO in male mice. Data are means ± SEM from 6 (DMSO) and 7 (L689) slices/mice. (**C**) Representative field responses obtained after LTP and after LFS delivery (overlayed on a sample baseline trace of reduced opacity) in the presence of DMSO (black) or L689 (grey). (**D**) Average percent cDEP in the presence of DMSO versus L689 (unpaired Student’s t test, mean difference = 12.6 ± 30.3%, p = 0.687).

**Fig. S2.**
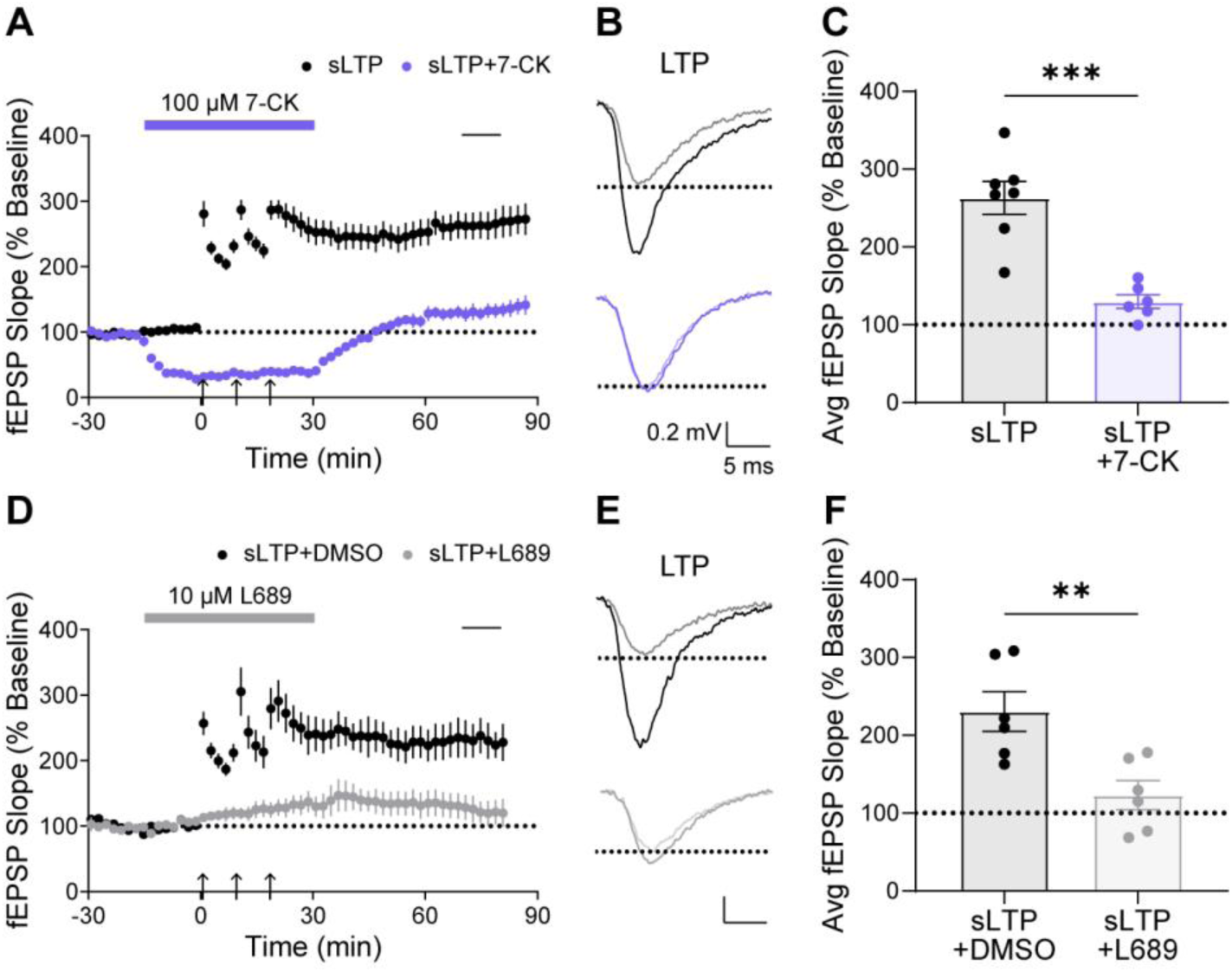
NMDAR glycine site-specific antagonists block sLTP. (**A**) 100 µM of 7-CK blocked sLTP induction (purple) ordinarily induced with a spaced TBS (black) in male mice. Data are means ± SEM from 7 (sLTP) and 6 (sLTP+7-CK) slices/mice. (**B**) Representative field responses obtained at baseline and after spaced TBS administration in control conditions (black) and in the presence of 7-CK (purple). (**C**) 10-minute average after LTP induction, indicated by the black bar in (A) in control conditions versus in the presence of 7-CK (unpaired Student’s t test, mean difference = 133.5 ± 24.3%, p = 0.0002). (**D**) 10 µM of L689 in DMSO blocked sLTP induction (grey) ordinarily induced with a spaced TBS in the presence of vehicle (0.01% DMSO) alone (black) in male mice. Data are means ± SEM from 6 slices/mice per group. (**E**) Representative field responses obtained at baseline and after spaced TBS administration in the presence of DMSO (black) or L689 (grey). (**F**) 10-minute average after LTP induction, indicated by the black bar in (D) in the presence of vehicle versus L689 (unpaired Student’s t test, mean difference = 107.6 ± 31.6%, p = 0.0067). All drugs were applied beginning 15 minutes before the first TBS administration and washed out 10 minutes after the final TBS administration, as indicated by the bars in (A) and (D). All scalebars are 5 ms x 0.2 mV.

**Fig. S3.**
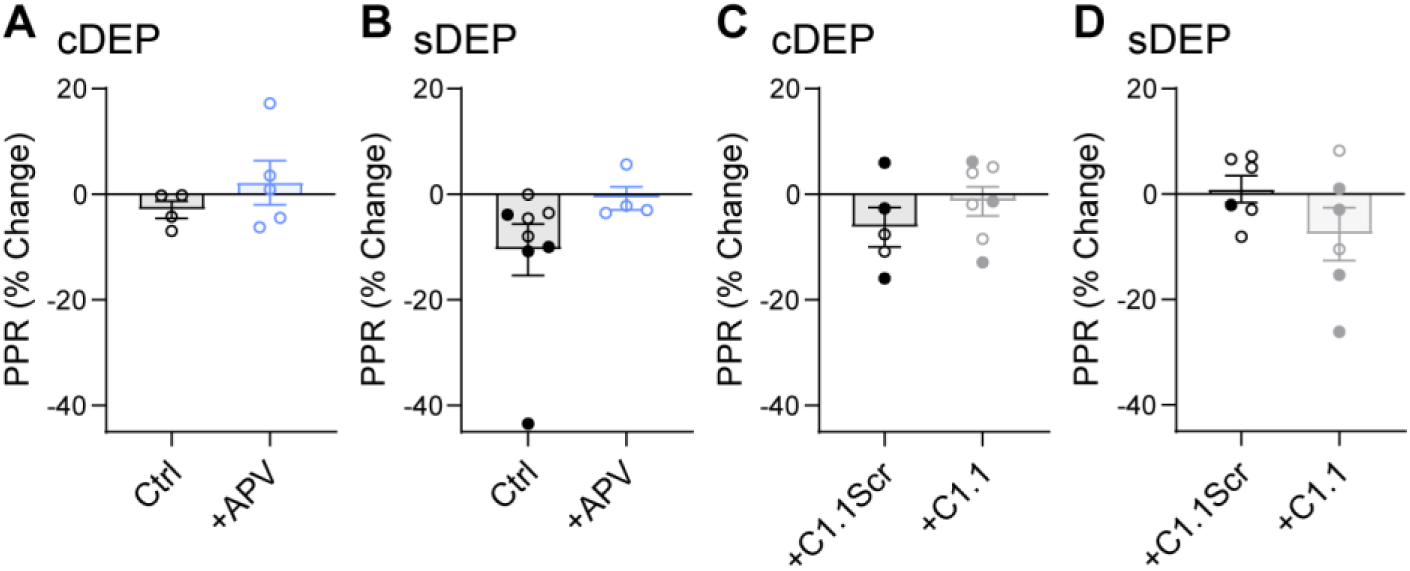
Paired pulse ratios (PPRs) as a percentage change between LTP and DEP induction are unaltered by drug treatments during DEP, consistent with a postsynaptic locus of expression. (**A**) The PPR percent change did not differ significantly in the absence or presence of NMDAR glutamate-site antagonist APV following cDEP induction (mean difference = −5.1 ± 4.9%, p = 0.333) or (**B**) sDEP induction (mean difference = −9.8 ± 7.2%, p = 0.206). Data are means ± SEM from 4 (cDEP), 5 (cDEP+APV), 8 (sDEP) and 4 (sDEP+APV) slices/mice. (**C**) The PPR percent change was not statistically significantly different in the presence of C1.1Scr versus C1.1 following cDEP induction (mean difference = −4.9 ± 4.5%, p = 0.303) or (**D**) sDEP induction (mean difference = 8.5 ± 5.6%, p = 0.160). Data are means ± SEM from 5 (cDEP+C1.1Scr), 7 (cDEP+C1.1), 6 (sDEP+C1.1Scr) and 6 (sDEP+C1.1) slices/mice. All statistical tests were unpaired Student’s t tests. Data from male (closed circles) and female (open circles) combined. Ctrl = ACSF only control.

**Fig. S4.**
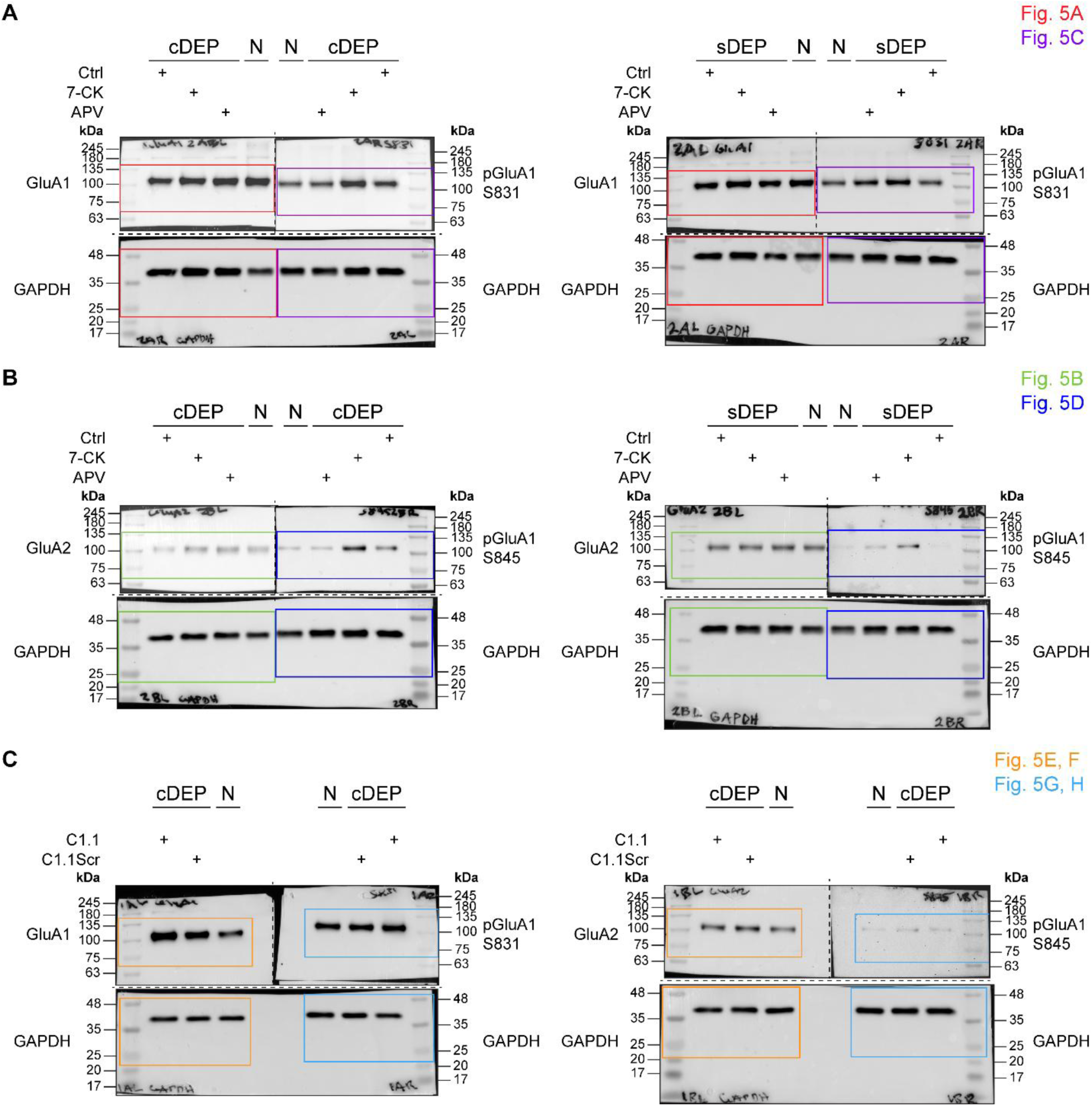
Full western blots corresponding to main Figure 5. (**A**) Full blots stained for GluA1 (top left-hand side of blots), pGluA1 S831 (top right-hand side of blots) and GAPDH (bottom section of blots) after membranes were cut as indicated along dotted lines. Blots containing samples frozen after cDEP (left) and sDEP (right) are shown. Samples were treated during electrophysiology experiments with 7-CK or APV as indicated. (**B**) Full blots stained for GluA2 (top left-hand side of blots), pGluA1 S845 (top right-hand side of blots) and GAPDH (bottom of blots) after blots were cut along dotted lines. Blots containing samples frozen after cDEP (left) and sDEP (right) are shown with corresponding treatments indicated. (**C**) Full blots stained for GluA1 (top left-hand side of left blot), pGluA1 S831 (top right-hand side of left blot), GluA2 (top left-hand side of right blot) and pGluA1 S845 (top right-hand side of right blot). Cropped blots from main Figure 5 are indicated with the corresponding colour-coded boxes. Images of blots stained for pGluA1 S831 and S845 were mirrored in Figure 5 for clarity.

